# Understanding Neuromodulation Pathways in tDCS: Brain Stem Recordings in Rat During Trigeminal Nerve Direct Current Stimulation

**DOI:** 10.1101/2023.09.14.557723

**Authors:** Alireza Majdi, Boateng Asamoah, Myles Mc Laughlin

**Author notes:** Correspondence Myles Mc Laughlin, Ph.D. Exp ORL, Department of Neuroscience, Leuven Brain Institute, KU Leuven, Leuven, Belgium. The data emerging from this research have been partly presented at 13^th^ FENS 2022, 1st BeNe Brain Stimulation 2022, and 5^th^ BRST 2023 Conferences in Paris, Hasselt, and Lisbon, respectively.

## Abstract

**Background:** Recent evidence suggests that transcranial direct current stimulation (tDCS) indirectly influences brain activity through cranial nerve pathways, particularly the trigeminal nerve. However, the electrophysiological effects of direct current (DC) stimulation on the trigeminal nerve (DC-TNS) and its impact on trigeminal nuclei remain unknown. These nuclei exert control over brainstem centers regulating neurotransmitter release, such as serotonin and norepinephrine, potentially affecting global brain activity.

**Objectives:** To investigate how DC-TNS impacts neuronal activity in the principal sensory nucleus (NVsnpr) and the mesencephalic nucleus of the trigeminal nerve (MeV).

**Methods:** Twenty male Sprague Dawley rats (n=10 each nucleus) were anesthetized with urethane. DC stimulation, ranging from 0.5 to 3 mA, targeted the trigeminal nerve’s marginal branch. Simultaneously, single-unit electrophysiological recordings were obtained using a 32-channel silicon probe, comprising three one-minute intervals: pre-stimulation, DC stimulation, and post-stimulation. Xylocaine was administered to block the trigeminal nerve as a control.

**Results:** DC-TNS significantly increased neuronal spiking activity in both NVsnpr and MeV, returning to baseline during the post-stimulation phase. When the trigeminal nerve was blocked with xylocaine, the robust 3 mA trigeminal nerve DC stimulation failed to induce increased spiking activity in the trigeminal nuclei.

**Conclusion:** Our results offer initial empirical support for trigeminal nuclei activity modulation via DC-TNS. This discovery supports the hypothesis that cranial nerve pathways may play a pivotal role in mediating tDCS effects, setting the stage for further exploration into the complex interplay between peripheral nerves and neural modulation techniques.

**Highlights:**

- Direct current stimulation of the trigeminal nerve (DC-TNS) modulates neural activity in rat NVsnpr and MeV.
- Xylocaine administration reversibly blocks the DC-TNS effect on neural responses.
- Trigeminal nerve stimulation should be considered a possible mechanism of action of tDCS.

## Introduction

Transcranial direct current stimulation (tDCS) is a non-invasive electrical stimulation method, in which electrodes are placed on the scalp to administer a weak direct electrical current (1 to 2 mA) [1]. tDCS has significant potential as a treatment option due to its affordability, portability, safety, and user-friendly nature [2]. However, understanding the underlying mechanisms of action remains a primary challenge in tDCS research [3].

One widely recognized effect of tDCS is its ability to alter membrane polarity in cortical neurons, thereby influencing the generation of action potentials [4, 5]. However, the view that current passes through the scalp, skull, and cerebrospinal fluid, to directly affect cortical neurons, is a matter of debate [6–8]. An alternative, somewhat controversial, explanation is that the impact of tDCS on neural circuits may also be indirect, acting through peripheral nerves such as the trigeminal and the greater occipital nerves [3, 6–8]. Recent evidence indicates that non-invasive transcutaneous direct current (DC) electrical stimulation of the greater occipital nerve establishes a ‘bottom-up’ communication pathway from the periphery via the brainstem (nucleus tractus solitarius (NTS)) and then the reticular formation to the relevant subcortical and cortical regions [6, 7].

Similarly, the trigeminal nerve has extensive connections to the brainstem and various other brain structures. It projects to the trigeminal nuclei, e.g., principal sensory nucleus (NVsnpr) and mesencephalic nucleus (MeV), which reciprocally project to locus coeruleus (LC), raphe nuclei (RN), and NTS [9, 10]. The LC and RN are rich in noradrenergic and serotoninergic innervation, respectively [9]. Additionally, the NTS is linked to glutamatergic and GABAergic systems [11]. Through these projections and connections, an increase in the activity of trigeminal nuclei can have several behavioral and physiological consequences [12] including enhanced vigilance [13, 14], accelerated mental information processing [15], reduced response delays [16], augmentation of the oxygen-dependent signal in cerebral blood flow, and mydriasis in response to tasks [17].

The NVsnpr is composed of neurons that are believed to be either sensory relay neurons, firing in proportion to stimulus intensity, or neurons that transform depolarization into bursts to alter sensory signals [18, 19]. Evidence also indicates that neurons within this nucleus can adjust their response stability based on specific stimulus attributes, such as duration and frequency [20]. Besides, it has been reported that trigeminal nerve stimulation elicits diverse activity patterns in MeV neurons [21].

The electrophysiological effects of DC trigeminal nerve stimulation (TN-DCS) on neuronal activity in the NVsnpr and MeV are little known. In this study, we aimed to investigate the effects of TN-DCS on neuronal activity within the NVsnpr and MeV in rats.

## Materials and methods

### Animals

For all experiments, we used 20 (n=10 for each nucleus) adult male Sprague Dawley rats (Charles River Laboratories, Belgium) weighing between 250-450 g at the start of each experiment. Before surgery, the rats were housed in pairs per cage. The animals were individually housed in cages in a temperature- and humidity-controlled room, maintaining a diurnal light-dark cycle with ad libitum access to food and water. All animal experiments adhered to the ARRIVE guidelines and were in accordance with the EU Directive 2010/63/EU for animal experiments. The experimental procedures conducted in this study received approval from the KU Leuven ethics committee for laboratory experimentation (License numbers: 201/2018 and 072/2020).

### Anesthesia and surgery

During the experiment days, the rats were anesthetized via an intraperitoneal (i.p.) injection of a 1.5 g/kg body weight dose of urethane (ethyl carbamate; Sigma-Aldrich, U2500) [22]. Subsequently, they were positioned in a stereotaxic frame (World Precision Instruments Ltd., Stevenage, UK) on a heated pad, while the core temperature was monitored using a metal rectal probe. The depth of anesthesia was regularly assessed by observing the toe-pinch reflex. Approximately 100 µL of urethane was administered via i.p. injection as needed to maintain a consistent anesthesia level. Once the scalp was removed, the skull was carefully adjusted to achieve a level of 0 degrees on both the anteroposterior and mediolateral planes. This adjustment ensured that the lambda-bregma and lateral ridges differences were less than 200 μm. At the end of each experiment, the animal was euthanized using an overdose i.p. injection of sodium pentobarbital (Dolethal ND, Vetoquinol, France) at 200 mg/kg [23].

### Stereotaxic coordinates and electrode placement

The Paxinos and Watson rat brain atlas was used to locate the NVsnpr relative to Bregma [24]. To this end, a burr hole craniotomy was performed using the 78001 Microdrill (RWD Life Science Co., Shenzhen, China) at the following coordinate AP: -9.0 mm, ML: ±2.9 mm, and DV:-7.5-9 mm. The MeV was located using the criteria described by Totah et al., [22]. A cranial window was made at AP: -4 to -5 mm and ML: ±1.2 mm relative to true lambda [24]. The dura was retracted and the electrode was lowered at a 15⁰ angle to the horizontal plane to avoid the transverse sinus overlaying the target area until reaching 5-7 mm below the cortical surface [22]. The electrode location in the MeV was confirmed by the audible presence of jaw movement-responsive cells [22].

### Stimulation protocol

To stimulate the trigeminal nerve’s marginal branch, a crocodile clip was positioned at the lower lip of the rat serving as an anode. Another clip was attached to the tail to serve as a cathode. The contact surfaces of the electrodes with the rat were covered with conductive gel (Signa Gel, Parker Labs, New Jersey). DC electrical stimulation was provided by an AM 2200 analog current source (AM Systems, Sequim, WA) connected to two bolt crocodile clips mentioned earlier. The current source received an analog voltage waveform generated from a data acquisition card (NI USB -6216, National Instruments, Austin, TX) operating at a sampling rate of 30 kHz. Dedicated MATLAB 2014 (MathWorks, Natick, MA, USA) controlled the data acquisition card. DC stimulation was delivered at ±0.5, ±1, ±2, and ±3 mA. The stimulation protocol consisted of the following sequence: 1 minute without any stimulation (pre), followed by 1 minute of DC stimulation (during), and finally another 1 minute without any stimulation (post).

In a control experiment, the trigeminal nerve was blocked by injecting 1 ml xylazine 2% (Xyl-M; VMD, Belgium) into the nerve and its branches on the face. This allowed us to test whether brain stem nuclei responses were due to activation of the trigeminal nerve or the result of volume conduction.

### Recording setup

We recorded neurons using a 50 μm thick silicone probe with 32 channels spanning across 1550 µm (Atlas Neuro, Leuven, Belgium, Model: E32+R-50-S1M-L20 NT). The output of the channels was amplified (x200) and digitized using a 16-bit, 30kHz acquisition board and 32-channel recording headstage (RHD 32 by Intan Technologies, Los Angeles, CA, USA). The probe data were monitored and recorded throughout the experiment on a PC hard drive at a sampling rate of 30 kHz via the Open Ephys GUI (https://open-ephys.org/).

### Data analysis

We performed spike sorting using SpyKING CIRCUS 1.0.1. [25]. Briefly, the data first underwent several preprocessing steps. SpyKING CIRCUS filtered all the signals with an auto-defined cut-off frequency for a Butterworth filter at 100 Hz (template width of 3 ms and a spatial radius of 250 µm for the templates). Then, spikes were automatically detected considering λ = 6 in the spike threshold (θ_k_) calculation formula. Accordingly, spatial whitening was done to eliminate erroneous spatial correlations among adjacent recording electrodes. Afterward, the dimensionality of the temporal waveforms was decreased by basis estimation (principal component analysis). Following the preprocessing steps, the data were clustered, and template matching as well as automated merging (spike trains binning) were performed [25]. The clustering was subsequently manually examined and adjusted using phy a GUI-based software for manual curation (https://github.com/cortex-lab/phy) and only distinct single-unit clusters were selected for further analysis. Our assessment of waveform quality involved detecting potential action potentials, and only clusters that showed recognizable action potential waveforms and had a V-shaped autocorrelogram were labeled as “good” before being exported to MATLAB. Following this, we proceeded to refine the data by checking for interspike interval (ISI) violations, which occurred when a putative unit fired more than twice within a 2-millisecond interval. Clusters that exhibited over 3% ISI violations were discarded from the analysis. Following the manual clustering phase, we further examined the distinct neuronal units in MATLAB. In MATLAB the mean spike rate in the 1-minute pre, 1-minute during, and 1-minute post conditions was calculated for further statistical analysis. Additionally, a spike rate over time waveform was calculated by taking the mean spike rate over 1-second bins.

### Statistics

The data processing and statistical analyses were conducted using MATLAB 2022a (MathWorks, Natick, MA, USA). In accordance with recommendations of [26], we employed a linear mixed-effect (LME) model to assess the impact of stimulation amplitude and condition (pre, during, and post) on spike rate for both NVsnpr and MeV data. We treated stimulation amplitude, condition, and their interaction as fixed effects while considering rats and neurons as random effects. The model codes in MATLAB’s fitlme.m were as follows:

1. Spike Rate ∼ Amplitude + Condition + Amplitude × Condition + (1|Neuron) + (1|Rat) to test the effect of amplitude and conditions on spike rate,
2. Spike Rate ∼ Amplitude + Condition+ Polarity + Amplitude × Condition × Polarity + (1|Neuron) + (1|Rat) to test the effect of polarity on spike rate, and
3. Spike Rate ∼ 1 + Xylo × Condition + (1 | Neuron) + (1 | Rat) to test the effect of xylocaine blockage on spike rate.

The ANOVA was then performed to test for the significance of the model’s fixed effects and interactions. To conduct post-hoc testing, we employed a two-sided Wilcoxon signed rank test with a strict Bonferroni correction to account for all possible multiple comparisons. For the NVsnpr dataset, this involved 4 stimulation amplitudes and 3 conditions, resulting in 12 multiple comparisons. We had 3 stimulation amplitudes and 3 conditions for the MeV dataset, leading to 9 multiple comparisons. The Akaike information criterion (AIC), Bayesian Information Criterion (BIC), and likelihood ratio tests were utilized to evaluate the optimal model fit for the data [27]. In all comparisons, a significance level of 0.05 was considered.

## Results

### Single unit response to TN-DCS in NVsnpr and MeV nuclei

Fig. 1A depicts a schematic diagram of the Sprague Dawley rat brain sagittal plane showing the relative position of stimulating and recording electrodes and the positional relationship of the NVsnpr, MeV, and trigeminal nerve. It also shows an example spike train from the NVsnpr (Fig. 1B) and MeV (Fig. 1C). TN-DCS is turned on at the 60-second time point and there is a noticeable increase in spike rate in both of the trigeminal nuclei in these examples. Individual spikes as identified after spike sorting are indicated with a red dot. Fig. 2 illustrates single unit responses in the NVsnpr (upper panel) and the MeV (lower panel) to three different stimulation amplitudes; represented by yellow, red, and blue lines for 3, 2, and 1 mA, respectively. This figure shows an example recording of the pre-during-post conditions during TN-DCS. that note that in the example shown, while the spike rate increases, the spike shape does not appear to be affected by the stimulation conditions.

**Fig. 1.**
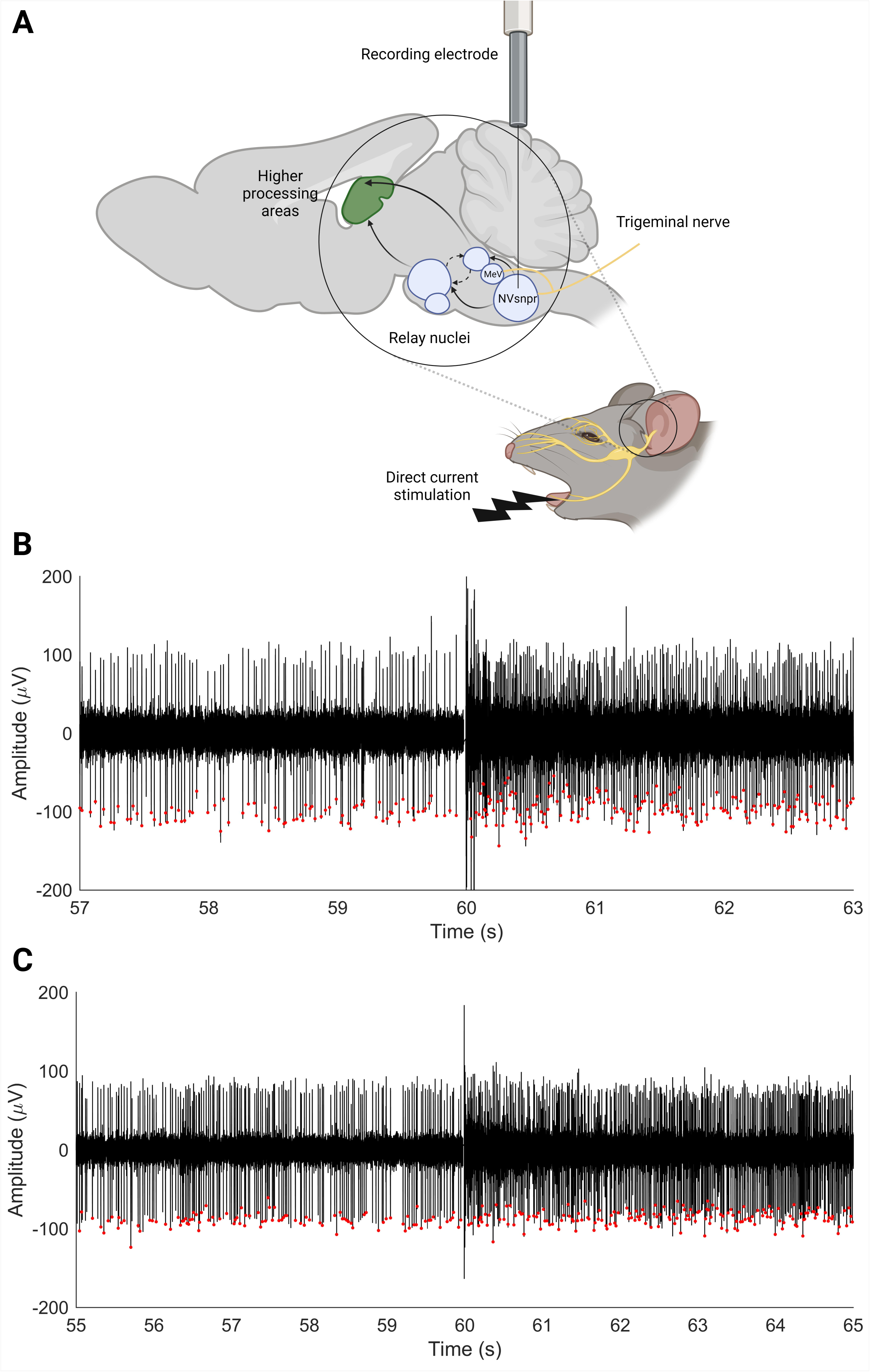
A) Schematic diagram showing stimulation and recording electrodes and the relative positional relationship of the recorded nuclei in the sagittal section of the Sprague Dawley rat brain. B) Example of spike trains showing typical fast irregular firing of neurons in B) the NVsnpr and C) the MeV before and after stimulation onset. A sharp increase in the number of spikes is observed upon the initiation of DC-TNS. Red dots represent spikes detected by the spike sorting process. NVsnpr, principal sensory nucleus; MeV, mesencephalic nucleus; DC-TNS, direct current trigeminal nerve stimulation. Figure 1A is created with BioRender.com.

**Fig. 2.**
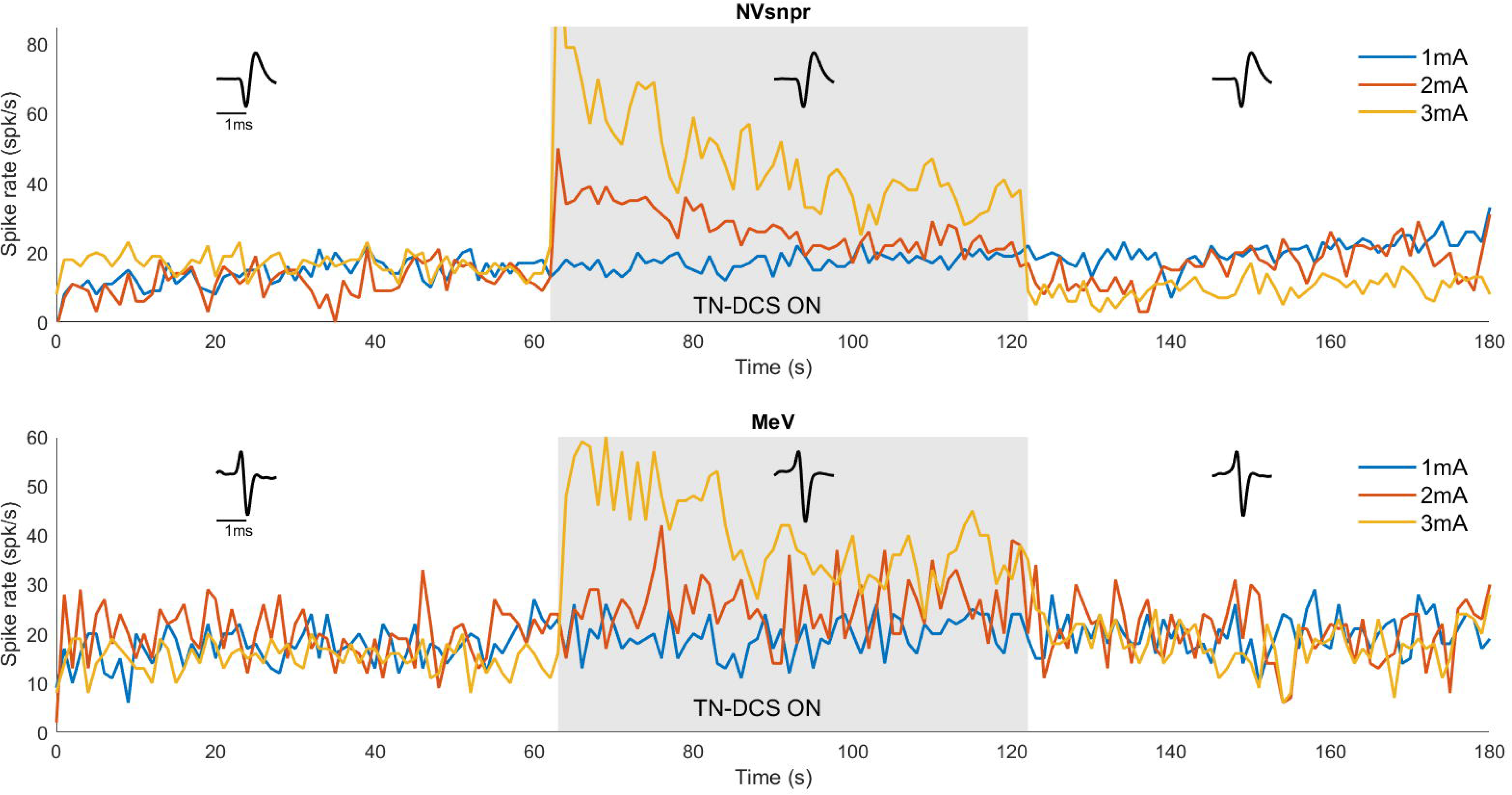
Individual neuron examples of the effect of 3 different amplitudes (blue, red, and yellow representing 1, 2, and 3 mA, respectively) of trigeminal nerve direct current stimulation (TN-DCS) on spike rate and shape in principle sensory nucleus (NVsnpr; top) and mesencephalic nucleus of trigeminal nerve (MeV; down) neurons. Application of 3 mA TN-DCS increases spike rate during stimulation in both nuclei without affecting spike shape. The y-axis represents the spike rate per second and the x-axis depicts time in seconds.

### Neuronal responses in NVsnpr to TN-DCS: Effect of stimulation amplitude, condition, and polarity

A total of 132 single units were isolated from the NVsnpr. Fig. 3 presents data for all these neurons grouped, illustrating their responses to TN-DCS under various stimulation amplitude ranges and polarities (Fig. 3A for anodic and Fig. 3B for cathodic amplitudes) across the pre, during, and post-conditions. The LME model was used to analyze the data.

**Fig. 3.**
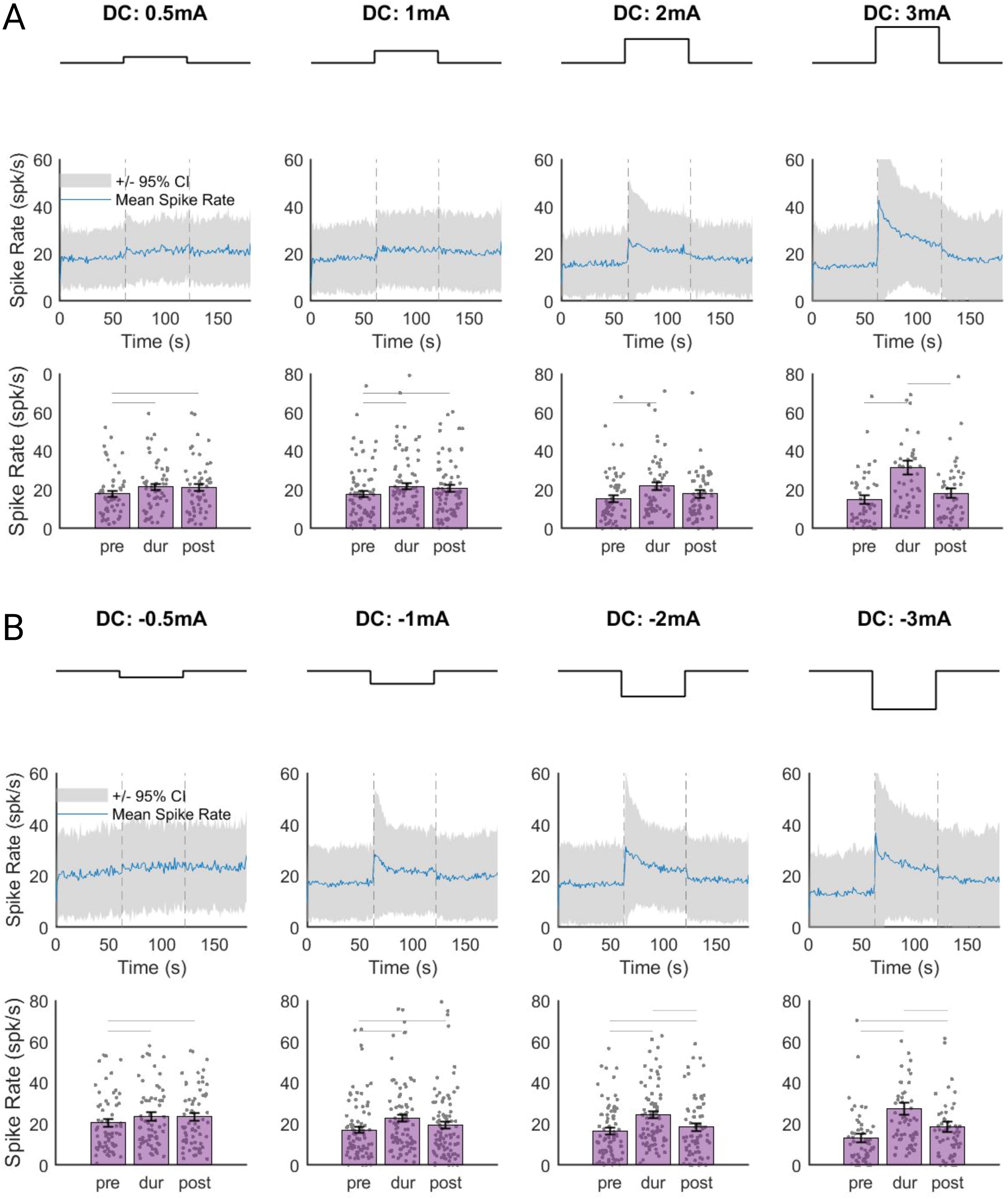
Effect of different amplitudes and conditions of trigeminal nerve direct current stimulation (TN-DCS) on neuronal spike rate in the principal sensory nucleus of the trigeminal nerve (NVsnpr). A) Anodic TN-DCS: The mean neuronal spike rate increased with increasing amplitude. The gray shadows and the blue lines (top) represent the 95% confidence intervals (CIs) and the average spike rate over time for all neurons. The bar graphs and error bars (down) display the mean spike rate and standard deviation (SD) for all pre, during, and post-TN-DCS conditions across different amplitude levels. A horizontal line denotes a significant increase in mean neuronal spike rate when switching from the pre-condition to the during-condition or from the during-condition to the post-condition. Each grey dot (down) represents a single unit. B) cathodic TN-DCS: The mean neuronal spike rate increased with increasing amplitude and remained significantly higher up to one minute after stimulation discontinuation. All figures in B follow the same style convention as in A. Data were analyzed using a linear mixed-effect model followed by a two-sided Wilcoxon signed rank test with a strict Bonferroni correction.

### Anodic stimulation

For the anodic stimulation, we found a significant impact of amplitude on spike rate (F_(1, 717)_ = 7.787, *p* = 0.005) and an interaction between amplitude and condition (F_(2, 717)_ = 7.951, *p* < 0.001). However, we found no significant effect of condition (F_(2, 717)_ = 1.073, *p* = 0.342) on the spike rate. This indicates that higher amplitudes of TN-DCS were associated with higher spike rates in the NVsnpr during stimulation (*p* < 0.001 for pre- vs. during stimulation conditions across all amplitudes, see Table 1 for full details of all post-hoc testing). Accordingly, higher anodic amplitudes were linked to higher Cohen’s d values indicating higher effect sizes (Table 1).

**Table 1.**
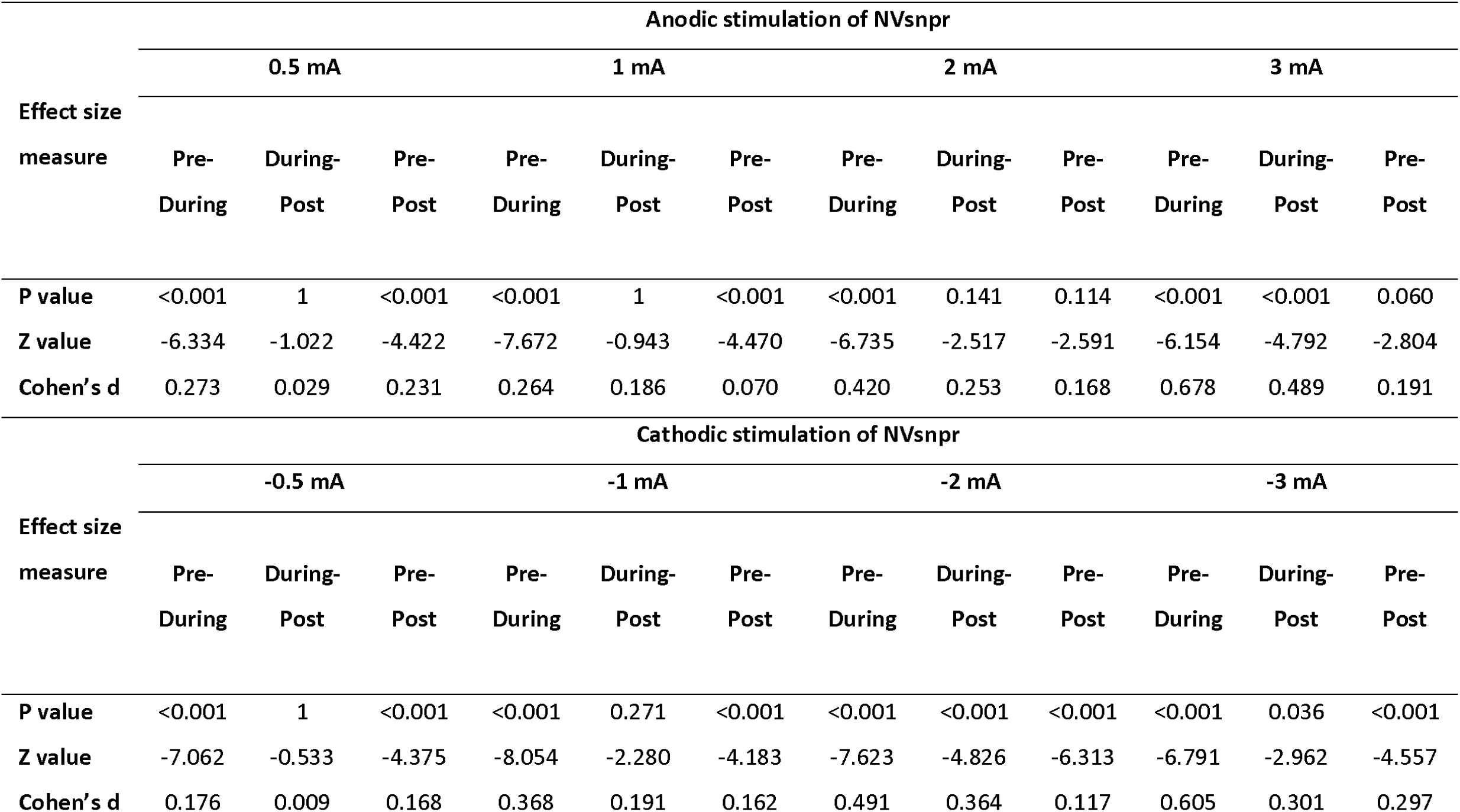
Size Post-hoc testing metrics for the impact of direct current trigeminal nerve stimulation (DC-TNS) on the spike rat in the principal sensory nucleus of the trigeminal nerve (NVsnpr) in rats. This table reports the effect size measures (Cohen’s d) along with p-values and Z-values (deviations from the mean) for different anodic and cathodic amplitudes of DC-TNS in rats under three conditions: pre- vs. during, post- vs. during, and pre- vs. post-stimulation.

### Cathodic stimulation

When analyzing the cathodic stimulation, we observed a significant effect of amplitude (F_(1,864)_ = 4.064, *p* = 0.044) and an interaction between amplitude and condition (F_(2, 864)_ = 4.058, *p* = 0.017), but there was no significant effect of condition alone (F_(2, 864)_ = 0.461, *p* = 0.630) on the spike rate. These results suggested that higher amplitudes of TN-DCS were linked to higher spike rates in the NVsnpr during compared to the pre-condition and interestingly, also in the post compared to the pre-stimulation condition (*p* < 0.001 pre- vs. during and pre- vs. post-stimulation conditions across all amplitudes, Table 1). Similar to the anodic condition, we found between increased cathodic stimulation amplitudes higher Cohen’s d values, indicating a larger effect size (see Table 1).

### Anodic vs. cathodic stimulation

We combined the datasets and added polarity as a fixed effect to test if there was a difference in the spike rate response to anodic and cathodic stimulation. Our analysis revealed no statistically significant effect of polarity on spike rates in the NVsnpr (F_(1, 1581)_ = 1.532, *p* = 0.215). With cathodic stimulation, the mean spike rate remained persistently high up to one minute after discontinuation of stimulation. However, this was not the case with anodic stimulation (Fig. 3).

### Neuronal responses in MeV to TN-DCS: Impact of amplitude and condition, and their interaction

A total of 74 single units were isolated from the MeV. Fig. 4 depicts the data for these neurons, grouped to illustrate their responses to TN-DCS across various amplitude ranges in the pre, during, and post-conditions. Our findings revealed a significant effect of amplitude (F_(1, 462)_ = 6.050, *p* = 0.014) and the interaction between amplitude and condition (F_(2, 462)_ = 3.367, *p* = 0.035) on the spike rate. However, there was no significant effect of the condition alone (F_(2, 462)_ = 0.402, *p* = 0.668) on the spike rate. This meant that higher amplitudes were associated with higher spike rates in the MeV in the during condition but not in the pre-or post-conditions (*p* < 0.001 for pre- vs. during stimulation conditions across all amplitudes, Table 2). The mean neuronal spike rate increased with increasing amplitude and remained significantly higher up to one minute after stimulation discontinuation (*p* < 0.001 for all comparisons). Similar to the NVsnpr, we found higher Cohen’s d values with higher anodic amplitudes, i.e., larger effect sizes (see Table 2 for full details).

**Fig. 4.**
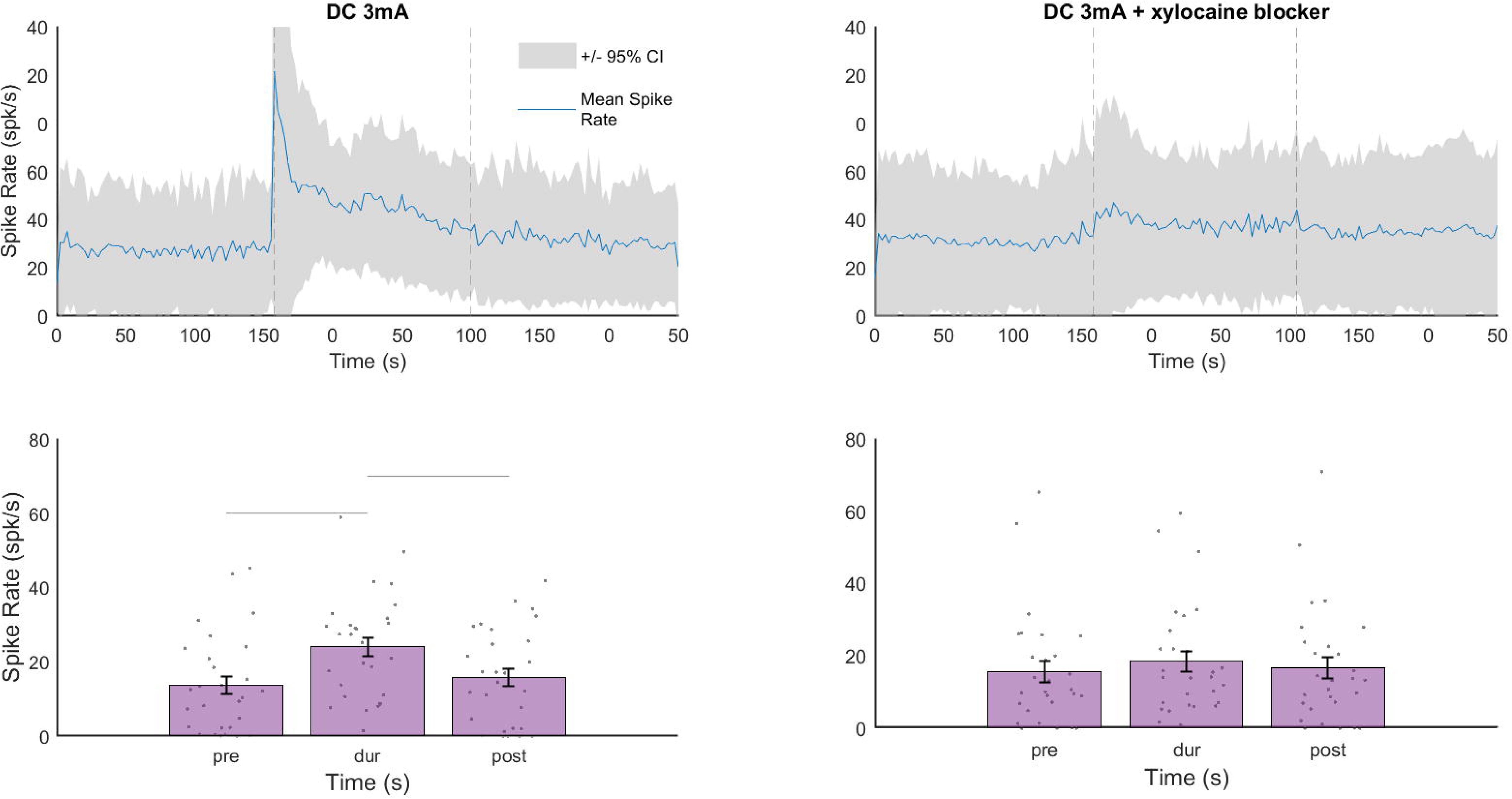
Effect of different amplitudes and conditions of trigeminal nerve direct current stimulation (TN-DCS) on neuronal spike rate in the mesencephalic nucleus (MeV) of the trigeminal nerve. Anodic TN-DCS: The mean neuronal spike rate increased with increasing amplitude. The gray shadows and the blue lines (top) represent the 95% confidence intervals (CIs) and the average spike rate over time for all neurons. The bar graphs and error bars (down) display the mean spike rate and standard deviation (SD) for all pre, during, and post-TN-DCS conditions across different amplitude levels. A horizontal line denotes a significant increase in mean neuronal spike rate when switching from the pre-condition to the during-condition or from the during-condition to the post-condition. Each grey dot (down) represents a single unit. Data were analyzed using a linear mixed-effect model followed by a two-sided Wilcoxon signed rank test with a strict Bonferroni correction.

**Table 2.**
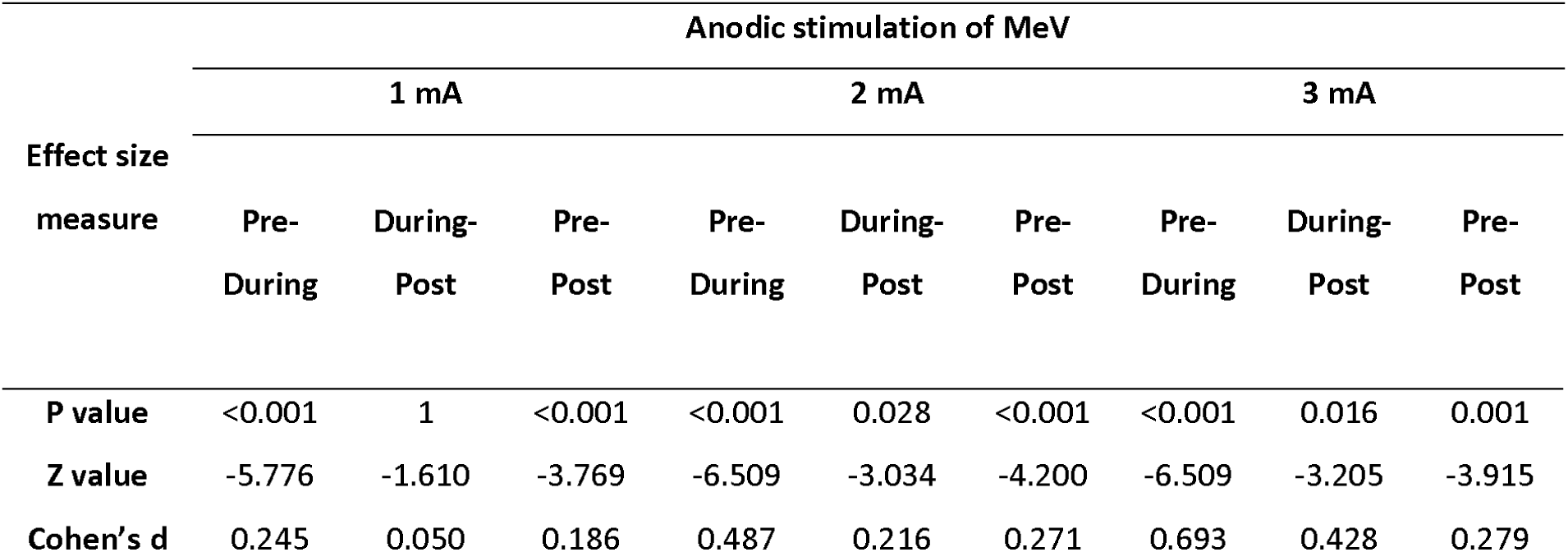
Effect size measures for the impact of direct current trigeminal nerve stimulation (DC-TNS) on the spike rat in the principal sensory nucleus of the mesencephalic nucleus (MeV) in rats. This table includes Cohen’s d values, p-values, and Z-values for various anodic amplitudes of DC-TNS in three conditions: pre- vs. during, post- vs. during, and pre- vs. post-stimulation. Higher absolute Z-values denote greater deviation from the mean, while larger Cohen’s d values signify more substantial effects.

### Neuronal responses in NVsnpr to TN-DCS: Effect of TN blockage via xylocaine

In the experiment with the xylocaine blocker, a group of 20 individual neurons (in a total of 5 animals) were isolated from the NVsnpr. The grouped data for these neurons are presented in Fig. 5, illustrating their responses to TN-DCS with and without xylocaine blocker. The results of our investigation revealed a significant effect of xylocaine on the spike rate in the NVsnpr (F_(1, 162)_ = 10.609, *p* = 0.001). Moreover, we observed significant effects of condition (F_(2, 162)_ = 12.855, *p* < 0.001) and interaction between xylocaine and the condition (F_(2, 162)_ = 3.521, *p* = 0.031) on the spike rate. This result indicates that TN blockage via xylocaine effectively inhibited the spike-rate effect of TN-DCS in the NVsnpr (*p* < 0.001 for pre vs. during condition before xylocaine injection vs. *p* > 0.05 for pre vs. during condition after xylocaine injection). See Table 3 for full post-hoc results.

**Fig. 5.** Effect of trigeminal nerve direct current stimulation (TN-DCS) at 3 mA amplitude on neurons’ spike rate in the trigeminal nerve’s principal sensory nucleus (NVsnpr) with and without xylocaine injection. A significant decrease in neuronal spike rate is observed after xylocaine injection (right panel). The gray shadows and the blue lines (top) represent the 95% confidence intervals (CIs) and the average spike rate over time for all neurons. The bar graphs and error bars (down) display the mean spike rate and standard deviation (SD) for all pre, during, and post-TN-DCS conditions across different amplitude levels. A horizontal line denotes a significant increase in mean neuronal spike rate when switching from the pre-condition to the during-condition or from the during-condition to the post-condition. Each grey dot (down) represents a single unit. Data were analyzed using a linear mixed-effect model followed by a two-sided Wilcoxon signed rank test with a strict Bonferroni correction.

**Table 3.** Effect size measures for the impact of direct current trigeminal nerve stimulation (DC-TNS) on the spike rat in the principal sensory nucleus of the mesencephalic nucleus (MeV) before and after unilateral xylocaine blocker injection to the marginal branch of the trigeminal nerve in rats. This table includes Cohen’s d values, p-values, and Z-values for 3 mA amplitude before and after blocker injection in three conditions: pre- vs. during, post- vs. during, and pre- vs. post-stimulation. Higher absolute Z-values denote greater deviation from the mean, while larger Cohen’s d values signify more substantial effects. NA, not applicable – no post-hoc testing was performed as there was no main effect.

| Anodic stimulation of NVsnpr |  |  |  |  |  |  |
| --- | --- | --- | --- | --- | --- | --- |
| Effect size measure | 3 mA without xylocaine blocker |  |  | 3 mA with xylocaine blocker |  |  |
|  | Pre-During | During-Post | Pre-Post | Pre-During | During-Post | Pre-Post |
| P value | <0.001 | 1 | <0.001 | 1 | 1 | 1 |
| Z value | -3.825 | -4.053 | -0.910 | NA | NA | NA |
| Cohen's d | 0.750 | 0.596 | 0.163 | NA | NA | NA |

## Discussion

In recent years, there has been a resurgence of enthusiasm surrounding tDCS, as it emerges as a promising instrument for modulating cortical function in the human brain and addressing neuropsychiatric disorders [28]. The appeal of tDCS lies in its capacity to non-invasively alter cortical activity and modulate excitability through undamaged cranial structures [28]. Numerous investigations have been undertaken in recent years to examine the physiological impacts and mechanisms of action of tDCS [29, 30]. There exists a consensus that the main mechanism by which DC stimulation influences the cerebral cortex is a slight alteration of the resting membrane potential of neurons (subthreshold depolarization or hyperpolarization depending on the polarity, duration, and strength of stimulation) [30, 31]. Recent research involving animals claims that tDCS, administered at intensities between 1 and 4.16 A/m², can exert a direct influence on subcortical structures, including the medial longitudinal fascicle and the red nucleus [32, 33]. The majority of the electrical current is attenuated by the scalp and skull, leaving only a small portion to reach the brain. Recent investigations propose that as much as 75% of the administered current does not effectively reach its target within the brain [34]. One potential rationale lies in the notion that tDCS might also impact neural circuits through an indirect route, namely, using peripheral nerves [5, 8].

Extensive empirical support substantiates the potential viability of trigeminal nerve stimulation (TNS) as a viable substitute for conventional therapeutic options in addressing neuropsychiatric disorders [35–39]. The clinical efficacy of TNS is well-established [40, 41]. However, the neurobiological mechanisms by which this efficacy is asserted remain largely unexplored [42].

The results of our study showed that TN-DCS stimulation of the trigeminal nerve with direct current significantly increased the mean spike rate in the NVsnpr during stimulation. With cathodic stimulation, the mean spike rate remained significantly high up to one minute after discontinuation of stimulation. However, this was not the case with anodic stimulation. Otherwise, we found no difference in the mean spike rate in the NVsnpr between the anodic and cathodic stimulations. Interestingly, these effects disappeared after administration of xylocaine to the marginal branch of TN where the stimulation electrode was positioned, suggesting that the increase in the mean spike rate in the NVsnpr was caused by direct stimulation of the trigeminal nerve and not any current spread to the nuclei under investigation.

In this study, the mean basic spike rate in the NVsnpr was found to be between 14.81 ± 2.26 Hz (mean ± standard deviation) (3 mA) and 18.01 ± 1.56 Hz (0.5 mA) which increased to 31.58 ± 3.43 Hz in 3 mA and 21.45 ± 1.69 in 0.5 mA, respectively. This is in line with the findings of Laturnus et al., who reported an increase in the basal median spike rate in the NVsnpr to 17.26 Hz [29.28 interquartile range (IQR)] after the introduction of the rat’s whiskers to 100 trials of seven-second white noise stimuli [43]. In humans, it has been shown that the application of acute, cyclic, 20-minute TNS leads to a noteworthy alteration in the activity of the bilateral polysynaptic pathway that involves areas such as the trigeminal nuclei in the brain stem [44].

We observed the same pattern in the MeV where the basic mean firing rate Increased from 16.98 ± 2.10 Hz to 20.82 ± 2.33 Hz in 1mA condition and from 15.72 ± 1.63 Hz to 26.63 ± 2.68 Hz in 3 mA condition. In an indirect relevance to this line of evidence, studies suggest that TNS via macrovibrissae movement causes an abrupt increase (with a very short latency of 1.3 ± 0.2 ms) in the neuronal discharge in the MeV which immediately returns to the basic level upon the termination of stimulation [45].

An increase in the activity of trigeminal nuclei can have several behavioral and physiological consequences [12–17]. There are several pathway mechanisms via which TNS has the potential to produce these effects, with one example pathway being vitae the LC. LC is a pivotal hub within the ARAS, serving as the primary origin of norepinephrine (NE) within the central nervous system [3]. There are multiple routes through which trigeminal input could reach both the ARAS and LC. It has been shown that each trigeminal nucleus, NVsnpr, and MeV included, sends projections to the LC [46–48]. Furthermore, trigeminal nerve stimulation could also reach the LC by traveling through the nucleus of the NTS and the reticular formation (RF) [49, 50]. These projections do not end in the LC core but in the periphery where the LC dendrites are located and thereby can influence the electrical coupling of LC neurons [51]. Interestingly, the decrease in trigeminal signals can result in lower levels of neurotrophic factors for LC neurons and thereby their dysfunction. This dysfunction can extend to the glial cells due to the strong electrical connection between MeV-LC cells and nearby astrocytes [52]. LC, a major noradrenergic area in the brain, can cause a desynchronization of neuronal activity, resulting in heightened levels of arousal and improved attention. This suggests that the LC plays a significant role in influencing these cognitive processes by modulating the electrical patterns in the brain. Furthermore, LC has the potential to impact an individual’s mood [17]. In addition, research has demonstrated that stimulating the LC can effectively reduce the perception of acute pain by reducing the release of neurotransmitters from nociceptive afferents. This may explain the analgesic effects rendered by TNS [53].

The trigeminal nerve, via its brain stim nuclei, also reaches NTS. Subsequently, this nucleus innervates various brain areas including LC, RN, and last but not least amygdala and hippocampus [54]. Moreover, the LC and the RN are connected through reciprocal pathways, indicating that these neuromodulatory regions are likely to interact when they are directly or indirectly activated by TNS [55–57].

These three brainstem regions i.e., LC, RN, and NTS, possess anatomical positioning that enables them to directly or indirectly impact the neurochemistry of extensive areas within the central nervous system. This may explain some of the behavioral and neurophysiological effects observed during TNS and tDCS [58].

To the best of our knowledge, this research represents a novel endeavor, being the first kind to characterize the effects of trigeminal nerve direct current stimulation on brainstem nuclei. The control of stimulation intensities and stimulation’s DC nature enabled an exploration of the dose-response relationship, providing valuable knowledge about the neural mechanisms at play during DC-TNS. Additionally, the use of extracellular recordings offered a high temporal resolution, ensuring the accurate measurement of spike rates and their dynamic changes in response to stimulation. However, several limitations should be acknowledged. First and foremost is the generalizability of the findings, as individual variations among animals may not be fully accounted for. Nevertheless, we tried to use the LME model to analyze our data and address this issue. Furthermore, the invasive nature of extracellular recordings raises concerns about potential alterations in the neural activity being studied, which may affect the reliability of the results. The limited duration of the 3-minute stimulation may not capture long-term effects or chronic responses adequately. It’s important to note that this study solely focuses on electrophysiological recordings, potentially overlooking other essential aspects of neural responses, such as changes in gene expression or synaptic plasticity. Lastly, the absence of behavioral assessments in the study limits our ability to gain a comprehensive understanding of the functional consequences of the observed neuronal changes. Interpreting the spike rate changes in these nuclei in response to electrical stimulation presents a complex challenge. Numerous factors can influence neuronal firing, including network effects and neurotransmitter interactions. Therefore, it is crucial to interpret the results within the broader context of neural function and connectivity.

## Conclusion

Our study demonstrated that both anodic and cathodic DC-TNS significantly increased neuronal spike rates in the NVsnpr and MeV nuclei during stimulation. These effects were reversible with xylocaine administration into the trigeminal nerve in rats. Contrary to the prevailing view, our findings suggest that peripheral nerve stimulation may play a substantial role in mediating the effects of tDCS, challenging the conventional notion of direct transcranial neuronal stimulation as the primary mechanism. However, it’s important to acknowledge the preliminary nature of this work. To provide a comprehensive understanding of the neural pathways involved, future investigations should encompass additional nuclei situated between the trigeminal nuclei and the target brain areas. This study serves as a starting point, encouraging further research and potential revisions of existing neuromodulation paradigms.

